# Ciliary ARL13B is essential for body weight regulation in mice

**DOI:** 10.1101/2023.08.02.551695

**Authors:** Tiffany T. Terry, Eduardo D. Gigante, Coralie M. Alexandre, Kathryn M. Brewer, Xinyu Yue, Nicolas F. Berbari, Christian Vaisse, Tamara Caspary

## Abstract

Primary cilia are sensory cellular appendages that regulate diverse developmental and homeostatic processes, including energy homeostasis. In animal models and humans, their dysfunction can lead to hyperphagia and obesity. ARL13B is a regulatory GTPase enriched in cilia. We engineered an *Arl13b* mouse allele, *Arl13b^V358A^*, that disrupts ARL13B from localizing to primary cilia. Homozygous *Arl13b^V358A/V358A^* mice become hyperphagic, obese, and insulin resistant. Restoring wildtype ARL13B to cilia in 4-week-old *Arl13b^V358A/V358A^* mice fully rescued the obesity and metabolic dysfunction. Additionally, the selective exclusion of ARL13B from cilia in the nervous system caused obesity. Together, these findings establish that ciliary ARL13B function within the nervous system is necessary for body weight regulation. Our ability to genetically uncouple the ciliary and non-ciliary functions of ARL13B in a cell-type-specific manner enables us to define its cilia-specific role and offers new insights into the molecular mechanisms underlying primary cilia control of energy homeostasis.

## INTRODUCTION

Primary cilia are small, nearly ubiquitous cellular appendages that act as signaling hubs on most cell types, including neurons involved in hunger, satiety, and energy homeostasis (1, 2). Disrupting cilia structure or the trafficking of specific receptors to these organelles leads to hyperphagia, obesity, and increased risk of diabetes. Human genetic data underscores this connection, where mutations in cilia-associated genes like *MC4R*, *ADCY3*, *INPP5E*, *ALMS1,* and the components of the BBSome cause severe early-onset obesity (1–3). *ALMS1* loss of function mutations lead to defects in cilia assembly and function, impacting ciliation and ciliary signaling in cells that regulate appetite and energy balance (4–6). BBS mutations disrupt cilia protein import and export, impacting ciliary signaling and trafficking of proteins involved in regulating food intake and energy metabolism (7–9). In adult mice, genetic ablation of cilia in specific neuronal cell types causes obesity due to hyperphagia and leads to elevated levels of leptin, glucose, and insulin (10, 11). While these findings show that primary cilia are required for regulating energy homeostasis, our inability to distinguish ciliary from non-ciliary functions of key cilia-associated proteins has limited the study of the molecular mechanisms by which cilia control energy homeostasis.

*Arl13b,* an ADP-ribosylation factor (ARF) protein family member, encodes a regulatory GTPase highly enriched on the ciliary membrane (12, 13). Like other regulatory GTPases, ARL13B has multiple functions, likely mediated by distinct effectors. For example, it has a conserved role as a guanine nucleotide exchange factor (GEF) for ARL3 (14, 15). In cilia, ARL13B retains the enzyme Inositol Polyphosphate-5-Phosphatase E (INPP5E), which regulates the phospholipid composition of the ciliary membrane (16, 17). In mice, loss of ARL13B is embryonic lethal, disrupting ciliogenesis and Hedgehog (Hh) signaling (18, 19). Patient mutations in *ARL13B* cause the ciliopathy Joubert Syndrome (JS) (20, 21). JS patients present with developmental delay, intellectual disability, and physical anomalies, and all known *ARL13B* JS-causing mutations disrupt ARL13B’s GEF activity for ARL3 (15).

We engineered an ARL13B mouse variant (ARL13B^V358A^) that maintains the known biochemical functions of ARL13B but disrupts ARL13B from localizing to primary cilia (22). This model overcomes a major challenge in studying ciliary signaling, as previous work often relied on genetic ablation of cilia, removing all ciliary signaling, or manipulating the signaling pathway in both the cilium and the cell. Thus, homozygous *Arl13b^V358A/V358A^* mice (hereafter called *Arl13b^A/A^*) reveal ciliary ARL13B function, after embryonic development and in adult animals.

Here, we report that *Arl13b^A/A^* mice became hyperphagic and obese, indicating that the cilia-specific function of ARL13B is critical for energy homeostasis regulation. ARL13B’s GEF activity for ARL3 is unlikely to be required to regulate body weight, as *Arl13b^R79Q^*mice expressing an ARL13B variant that lacks ARL13B GEF activity for ARL3 were not overweight. Introducing ciliary ARL13B in 4-week-old *Arl13b^A/A^* mice prevented obesity, suggesting ARL13B controls body weight through a homeostatic role in ciliary signaling. To control the subcellular localization of ARL13B in a tissue-specific manner, we developed an original approach, based on the combination of the *Arl13b^A^* and *Arl13b^floxed^* conditional alleles. Combining these alleles with *nestin-Cre* to specifically exclude ARL13B from cilia only in cells within the nervous system, we find that loss of ciliary ARL13B in the nervous system alters energy homeostasis. The cilia-specific exclusion of ARL13B provides a molecular entry point for understanding the role of cilia-mediated signaling in obesity, which will likely uncover mechanisms driving common forms of obesity.

## RESEARCH DESIGN AND METHODS

### Ethics Statement

All mice were cared for in accordance with NIH guidelines, and animal experimental procedures were approved by the Institutional Animal Care and Use Committees (IACUC) at Emory University (20170058), the University of California, San Francisco (AN201856), and Indiana University-Indianapolis (SC358R).

### Mouse lines

Mice were housed in a barrier facility and maintained on a 12:12 light cycle at an ambient temperature of 23°C ± 2°C and relative humidity of 50%–70%. Mice were fed with rodent diet 5058 (Lab Diet) and group housed (up to 5 mice per cage). Lines used were *Arl13b^V358A^* (C57BL/6J-*Arl13b^em1Tc^*) [MGI: 6256969] (22), *Arl13b^R79Q^* (C57BL/6J-*Arl13b^em2Tc^*) [MGI: 6279301] (23), *Arl13b^flox^(* ^tm1.1Tc^) [RRID: IMSR_JAX:031077]*), Arl13b-Fucci2a* [RRID: IMSR_EM:12168], *CAGGCre-ER^TM^ (Tg(CAG-cre/Esr1*)5Amc)* [RRID: IMSR_JAX:004682], *nestin-Cre (Tg(Nes-cre)1Kln* [RRID: IMSR_JAX:00377].

### Genotyping primers

Mice were genotyped for the V358A mutation using primers Forward: 5’-CAGTAAGAAGAAAACCAAGAAACTAAGACTCCTTTTCATTCATCGGGC-3’ and Reverse: 5’-GACAGTAAAGGATTCTTCCTCACAACCTGAC-3’ to detect the mutant allele and primers Forward: 5’-CTTAAGATGACTTTGAGTTTGGAAGAAATACAAGATAGC-3’ and Reverse: 5’-GCGTGGGACTCTTTGGAGTAGACTAGTCAATACAGACGGGTTCTA-3’ to detect the wildtype allele. Mice were genotyped for the R79Q mutation using primers Forward: 5’-CCAAGTTACCATCTTTGACTTAGGAGGTGG GAATTCTTTTTAGGGCATA- 3’ and Reverse: 5’-ACAGCAGCATCCCTGATACTTACACC-3’. Mice were genotyped for the *Arl13b^flox^*allele using primers Forward: 5’-AGGACGGTTGAGAACCACTG-3’ and Reverse: 5’-CGACCATCACAAGTGTCACC-3’. Mice were genotyped for the *Arl13b-Fucci2a* allele using primers Forward: 5’-AAAACCTCCCACACCTCCC-3’ and Reverse: 5’-CGACCATCACAAGTGTCACC-3’ to detect the transgene and primers Forward: 5’-AGGGAGCTGCAGTGGAGTAG-3’ and Reverse: 5’-CTTTAAGCCTGCCCAGAAGA-3’ to detect the wildtype allele. Mice were genotyped for *CAGGCre-ER^TM^* using primers Forward: 5’-CTCTAGAGCCTCTGCTAACC-3’ and Reverse: 5’-CCTGGCGATCCCTGAACATGTCC-3’. Mice were genotyped for *nestin-Cre* using primers Forward: 5’-CGCCGCTACTTCTTTTCAAC-3’ and Reverse: 5’-AATCGCGAACATCTTCAGGT-3’.

### Mouse metabolism studies

Mice were single-housed for 7 days, and food was weighed daily for at least 4 days. The average daily food weight was measured between groups. Energy expenditure was measured by the Comprehensive Lab Animal Monitoring System (CLAMS, Columbus Instruments, Columbus, Ohio). Mice were tested over 96 continuous hours, and the data from the last 48 hours were analyzed. Kilocalories per hour were calculated using the Lusk equation: Energy Expenditure = (3.815 + 1.232 × respiratory exchange ratio [RER]) × oxygen consumption rate (VO_2_) and analyzed with CalR software version 2 (24) (ANCOVA with lean mass used as a covariate). RER=VCO_2_/VO_2_. Lean mass and fat mass were measured using the EchoMRI^TM^ system (25).

### Glucose tolerance test

Adult mice were fasted for 16 hours and injected intraperitoneally with 1 g/kg body weight of glucose (Sigma G7021). Blood glucose was measured at 10, 20, 30, 60, 90, and 120 minutes via tail sampling with the AlphaTrak3 blood glucose meter (Zoetis). A blood sample for fasting glucose levels was taken at time point 0, before glucose was injected.

### Insulin tolerance test

Adult mice were fasted for 3 hours before intraperitoneal injection of 0.75 U/kg body weight of regular human insulin (Humulin R, Lilly USA; Novolin R, Novo Nordisk). Blood glucose levels were measured at 15, 30, 45, and 60 minutes. A blood sample for fasting insulin levels was taken at time point 0, before insulin was administered.

### Blood serum analysis

Blood was allowed to coagulate at room temperature for 30 minutes, then centrifuged at 1200 x g for 10 min at room temperature. Serum was collected and stored at - 80°C. Insulin and leptin levels were measured using Crystal Chem’s ultra-sensitive mouse ELISA kits according to the manufacturer’s protocol (leptin: catalog # 90080; insulin: catalog # 90030).

### Tamoxifen administration

Tamoxifen (Sigma T5648) stock solution was prepared at a concentration of 20mg/ml in 100% EtOH and stored at -20°C. Each dose of tamoxifen was freshly prepared in corn oil on the day of injection and dissolved using a speed vacuum centrifuge (Eppendorf Vacufuge plus). To induce gene expression, 10ul/g tamoxifen in 200ul corn oil was administered by oral gavage to 4-week-old mice using a 1ml syringe and a 22G 1.5-inch straight needle (Braintree Scientific Inc).

### Tissue harvesting and preparation

Mice were euthanized with isoflurane inhalation followed by perfusion with ice-cold PBS and ice-cold 4% paraformaldehyde (PFA). Brain and pancreas tissues were harvested and post-fixed with 4% PFA overnight at 4°C. Tissues were then cryoprotected in 30% sucrose in 0.1 M phosphate buffer at 4°C until tissues sank in solution. Samples were then frozen in optimal cutting temperature compound (Tissue-Tek OCT, Sakura Finetek).

### Immunohistochemistry

12µm or 20µm cryosections were rehydrated, blocked and permeabilized in antibody wash (5% heat-inactivated goat serum, 0.1% Triton X-100 in Tris-buffered Saline) for 30-40 minutes. Tissues were incubated with primary antibodies overnight at 4°C, washed three times with 0.1% Triton X-100 in Tris-Buffered Saline (TBST), and incubated with secondary antibodies for 1 hour at room temperature. Tissues were washed three times with TBST and incubated with Hoechst 33342 for 5 minutes. Slides were coverslipped with ProLong Gold (ThermoFisher) mounting media. Slides cured overnight at room temperature in the dark and were stored long-term at -20°C. Slides were imaged on a BioTek Lionheart FX microscope. Primary antibodies used were: mouse anti-ARL13B (1:1000, NeuroMab, N295B/66); rat anti-ARL13B (1:500, BiCell, 90413) rabbit anti-acetylated α-tubulin (1:1000,Cell Signaling, 5335); mouse anti-acetylated α-tubulin (1:2500, Sigma, T7451); chicken anti-ACIII (1:1000, Encor, CPCA-ACIII); rat anti-insulin (1:1000, R&D systems, MAB1417); rabbit anti-glucagon (1:2000, Abcam, ab92517), chicken anti-GFP (1:8000, Abcam, ab13970) recognizes cerulean. Secondary antibodies used were: goat anti-mouse AlexaFluor 488, donkey anti-rat AlexaFluor 488, goat anti-chicken AlexaFluor 647, goat anti-chicken AlexaFluor 488, goat anti-rat AlexaFluor 568 and donkey anti-rabbit AlexaFluor 555 (all at 1:500, ThermoFisher).

### AgRP and POMC neuron axonal projections

Mice were fasted for 48 hours prior to perfusion to increase AGRP levels in AgRP neuron projections. Mice were injected with leptin (R&D 498-OB, 5μg/injection/mouse administered twice daily at 7am and 7pm) for 3 days prior to perfusion to increase αMSH levels in POMC neuron projections. Brains were cryosectioned into 35 μm-thick free-floating sections. Three PVN sections (Bregma = -0.72mm, -0.8mm, or -0.94mm) from 48 hours fasted and leptin-injected mice were stained for AGRP and αMSH, respectively. To detect αMSH, tissues were unmasked in 0.3% glycine and 0.3% SDS for 10minutes each. Tissues were then blocked in 90% antibody buffer (1.125% NaCl, 0.75% Tris base, 1% BSA, 1.8 % L-Lysine, and 0.04% sodium azide; pH adjusted to 7.4), 10% donkey serum (Jackson ImmunoResearch, 017-000-121), and 0.3% Triton X-100. Tissues were incubated with the primary antibody at 4°C for 40 hours to detect AGRP, or overnight to detect αMSH. Tissues were mounted with Prolong Diamond Antifade Mounting Media (Invitrogen, P36970). Primary antibodies used were goat anti AGRP (1:1000, Neuromics, GT15023) and sheep anti-αMSH (1:500, Sigma, AB5087). Secondary antibodies used were donkey anti-goat AlexaFluor 488 and donkey anti-sheep AlexaFluor 488 (at 1:500 dilution). Images were acquired with a 20x objective on a Nikon Ti inverted fluorescence microscope equipped with a CSU-W1 wide-field-of-view confocal. AGRP and αMSH signals were segmented on Imaris 9.5.1 using the surface tool.

### RNAScope

Brains from 8-week-old adult mice were harvested, fixed, and prepared for RNAscope in situ hybridization according to the manufacturer’s protocols. Briefly, 15μm cryosections were mounted on slides and then post-fixed with 4% PFA for 16hr at 4 C. Detection of transcripts was performed using the RNAscope 2.5 HD Duplex Assay (Advanced Cell Diagnostics (ACD), Newark, CA). Tissue pretreatment was performed according to technical note 320534 Rev A. Probe hybridization, counterstaining, and mounting of slides were performed according to user manual no. 322500-USM Rev A. Slides were assayed using probes to AgRP (Cat No. 400711) and POMC (Cat No. 314081) transcripts (ACD). Sections were counterstained with hematoxylin, dehydrated, and mounted using VectaMount (Vectorlabs, Burlingame, CA). Slides with positive control probe (PPIB-C1/POLR2A-C2; Cat No. 321651) and negative control probe (DapB; Cat No. 320751) were run with each experiment. (n ≥ 3 animals per group).

### Data and Resources

All data generated or analyzed during this study are included in the published article (and its online supplementary files). No applicable resources were generated or analyzed during the current study.

## RESULTS

### Arl13b^A/A^ mice are obese, hyperphagic, and insulin-resistant

Complete ablation of ARL13B protein in mice (*Arl13b^hnn/hnn^* null mice) causes embryonic lethality; however, homozygous *Arl13b^A/A^*mice survive into adulthood and display an increased body weight compared to wildtype (*Arl13b^+/+^*) littermates (18, 22). To further characterize their body weight profile, we generated longitudinal data from weaning (week 3) to adult (week 10). We found no significant differences in body weight curves among the control genotypes: *Arl13b^+/+^*, *Arl13b^A/+^*, and *Arl13b^hnn/+^* in male or female mice, indicating the mutations are recessive and display no evidence of neomorphic (e.g. dominant negative) effect. The *Arl13b^A/A^* mice became significantly heavier than all control genotypes at week 5 for males and week 7 for females (**Figure 1A** and **B**). By week 10, both male and female *Arl13b^A/A^* mice were, on average, 33% heavier than controls.

**Figure 1:**
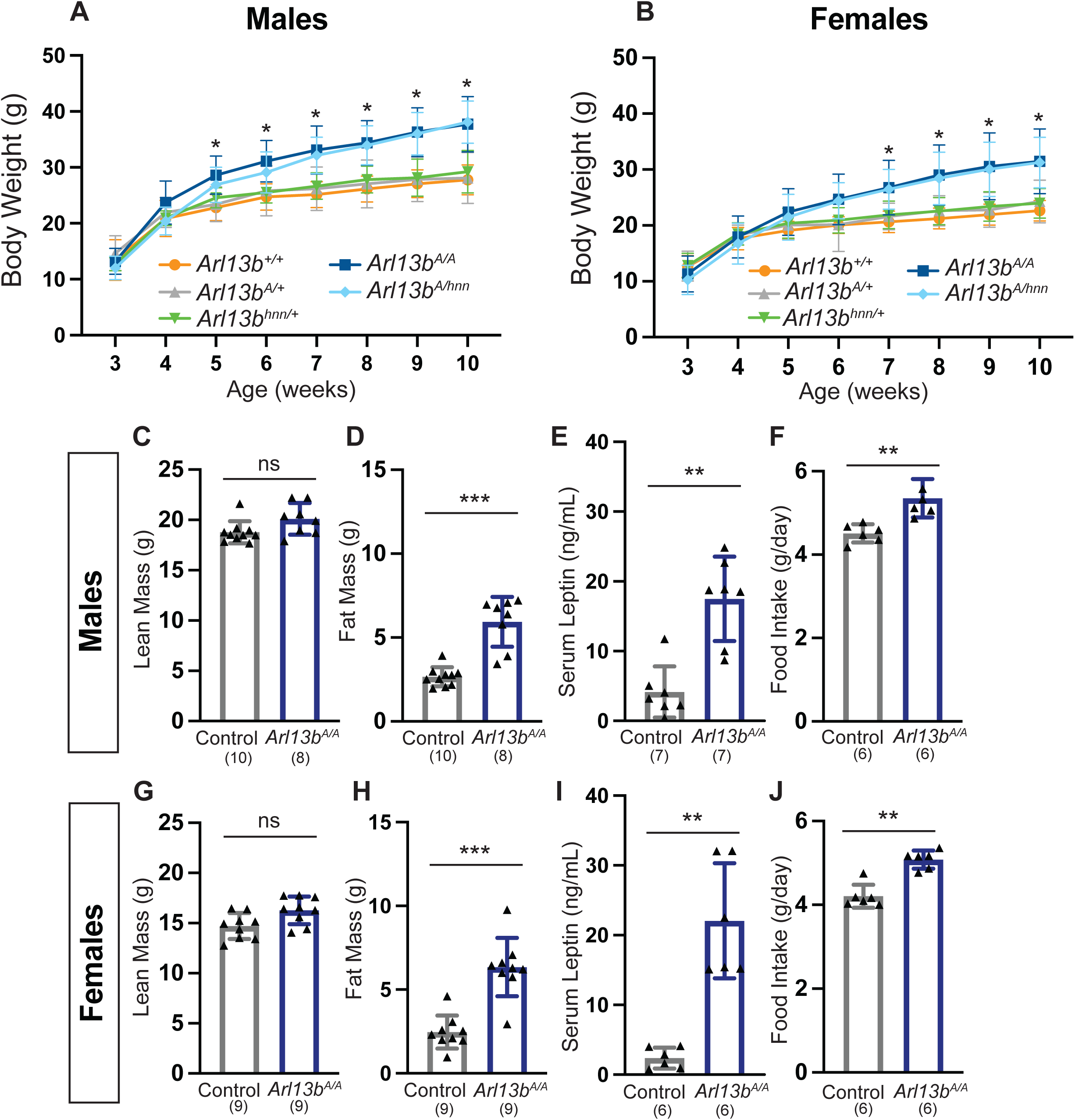
Exclusion of ARL13B from cilia leads to obesity. **(A-B**) Weekly male and female body weights from 3 to 10 weeks of age: *Arl13b^+/+^* (males n=14; females n=13), *Arl13b^A/+^* (males n=22; females n=22), *Arl13^hnn/+^* (males n=10; females n=9), *Arl13b^A/A^*(males n=24; females n=26), and *Arl13b^A/hnn^* (males n=15; females n=17). Statistical significance was determined using repeated-measures ANOVA followed by Tukey’s multiple comparisons test. Asterisk (*) indicates *Arl13b^A/A^*is significantly different (*P* < 0.05) from control (*Arl13b^+/+^*). (**C** and **G**) Lean mass and (**D** and **H**) fat mass of male and female control (*Arl13b^A/+^*) and *Arl13b^A/A^* (performed in mice 6 weeks old). (**E** and **I**) Serum leptin levels (analyzed in mice 10 weeks old). (**F** and **J**) Food intake (performed in mice 4-5 weeks old). Data are presented as means ± SD. Sample size indicated on graphs. Data in **C-J** were analyzed using the Mann-Whitney U test. \*\**P* < 0.01; \*\*\**P* < 0.001; ns, not significant with *P* > 0.05.

*Arl13b^A/A^* mice are obese, having an increase in fat mass. Body composition analysis showed comparable lean mass between *Arl13b^A/A^* and control animals (**Figure 1C** and **G**). The difference in body weight between the two genotypes was characterized by a 123% increase in fat mass in male *Arl13b^A/A^*mice and a 158% increase in fat mass in female *Arl13b^A/A^* mice (**Figure 1D** and **H**). Consistent with the increase in fat mass, we also observed increased leptin levels in *ad libitum*-fed *Arl13b^A/A^* mice (**Figure 1E** and **I)**.

To evaluate whether the *Arl13b^A/A^* weight phenotype is sensitive to gene copy number, we bred the *Arl13b^A^* allele with the null *Arl13b^hnn^* allele. We weighed male and female *Arl13b^A/hnn^* heterozygous mice from weeks 3 to 10. We found that *Arl13b^A/hnn^* mice displayed a similar body weight phenotype to *Arl13b^A/A^* mice, indicating that less overall ARL13B protein had no impact on this phenotype (**Figure 1A** and **B**).

To examine whether the obesity phenotype is the result of changes in feeding, we measured food intake from weeks 4 to 5, just prior to when we first observed an increase in *Arl13b^A/A^* body weight. We found that male and female *Arl13b^A/A^* mice consumed, on average, ∼20% more food per day than their littermate controls (**Figures 1F** and **J**). To investigate whether changes in activity and/or metabolism contribute to the increased body weight in *Arl13b^A/A^* mice, we measured energy expenditure, physical activity, and respiratory exchange ratio using a Comprehensive Lab Animal Monitoring System (CLAMS). Male *Arl13b^A/A^*mice displayed a modest 13% increase in daily energy expenditure, specifically during the light phase (**Supplemental Figure 1 A-C)**. Females showed no changes in energy expenditure compared to control animals (**Supplemental Figure 1 H-J**). We found no difference in the total physical activity and the total respiratory exchange ratio in male and female *Arl13b^A/A^* mice compared to control littermates (**Supplemental Figure 1 D-G and K-N**). These data demonstrate that the obesity we observe in male and female *Arl13b^A/A^* mice is due to hyperphagia.

We further determined if excluding ARL13B from cilia influenced glycemic regulation. *Arl13b^A/A^* mice were hyperinsulinemic (**Figure 2A** and **B**) and normoglycemic (**Figure 2 C-F**) at baseline, suggesting conserved insulin production by beta-cells. We performed glucose and insulin tolerance tests in mice aged 11-14 weeks. We found no significant differences in body weight among the control genotypes (*Arl13b^+/+^, Arl13b^A/+^, and Arl13b^hnn/+^*); therefore, we pooled data from these mice as a single control group. During glucose tolerance tests, both male and female *Arl13b^A/A^*mice exhibited elevated blood glucose levels (**Figure 2C** and **D**). Insulin did not lower blood glucose levels in male and female *Arl13b^A/A^* mice compared to controls, revealing an insulin-resistant phenotype consistent with the changes we observed in body composition (**Figure 2E** and **F**). Together, these data indicate that *Arl13b^A/A^* mice are hyperinsulinemic, glucose-intolerant, and insulin-resistant, congruent with their obese phenotype.

**Figure 2:**
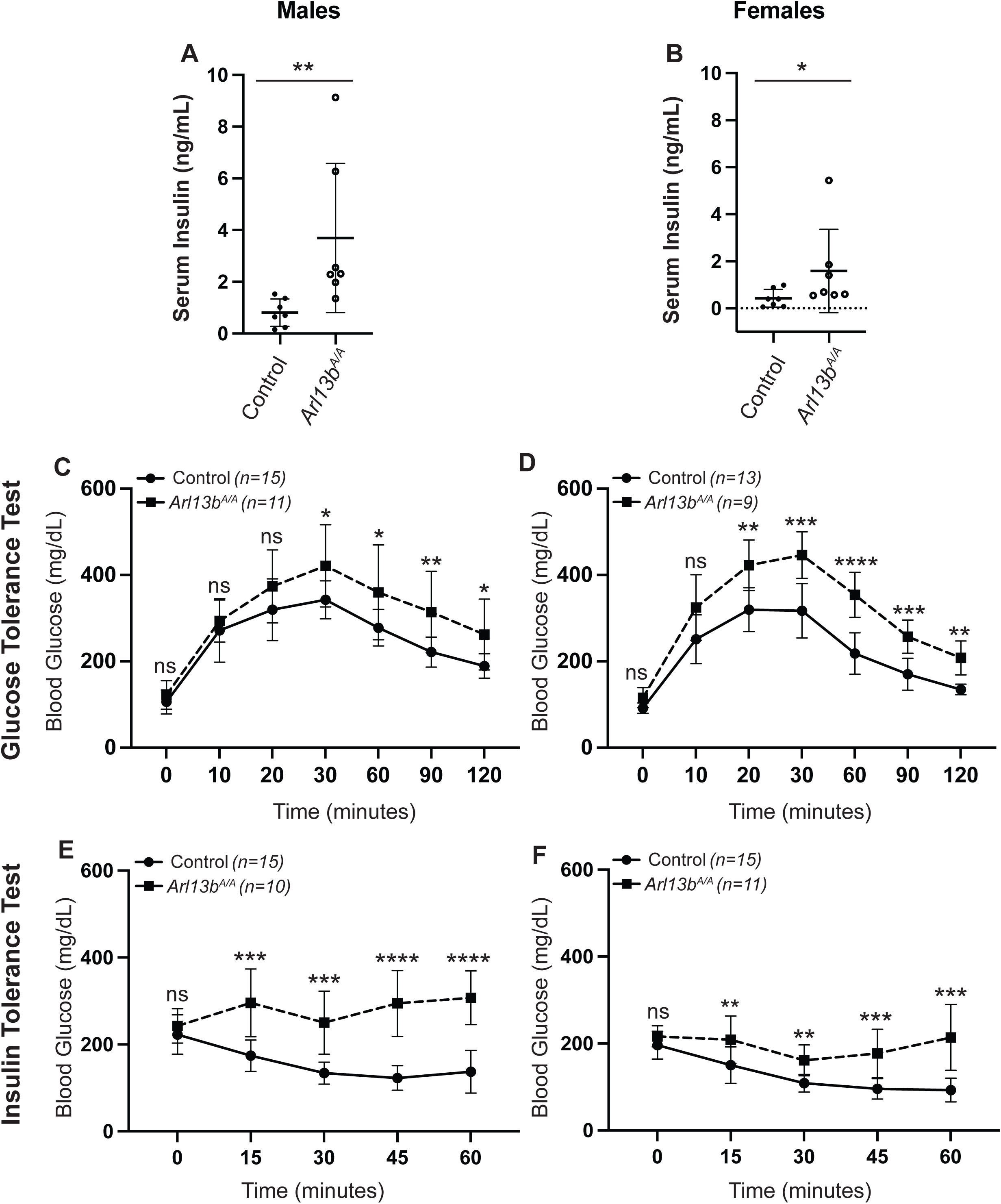
Insulin resistance and impaired glucose metabolism in *Arl13b^A/A^* mice. (**A-B**) Serum insulin levels in nonfasted 10-week-old male and female mice. Data points represent individual mice. Data are presented as means ± SD. Serum levels were analyzed using the Mann-Whitney U test. (**C**-**D**) Glucose tolerance test in males and females. Blood glucose was measured at the indicated times after i.p. glucose injection. (**E**-**F**) Insulin tolerance test of males and females. Blood glucose was measured at the indicated times after i.p. insulin injection. The control group comprised pooled data from *Arl13b^+/+^*, *Arl13b^A/+^*, and *Arl13b^hnn/+^* mice, as these mice did not differ in body weight. Data are presented as means ± SD. Sample size indicated on graphs. Statistical significance was determined using repeated-measures ANOVA followed by Fisher’s least significant difference (LSD) test. \**P* < 0.05; \*\**P* < 0.01; ns, not significant with *P* > 0.05.

### ARL13B^V358A^ protein is absent from cilia of tissues associated with ciliary control of energy homeostasis

Cilia are implicated in neuronal cell types in the hypothalamic nuclei for their roles in regulating feeding and energy homeostasis (1). They are also linked to insulin release in the pancreas (26). The engineered ARL13B^V358A^variant disrupts ARL13B from localizing to primary cilia in mouse embryonic fibroblasts, neural tube, and kidneys (22, 27, 28) (**Figure 3A** and **B**). To validate that ARL13B^V358A^ protein is also excluded from cilia of cell types relevant to metabolic control, we performed immunofluorescence staining for ARL13B in the endocrine pancreas and hypothalamus of *Arl13b^A/A^* mice.

**Figure 3:**
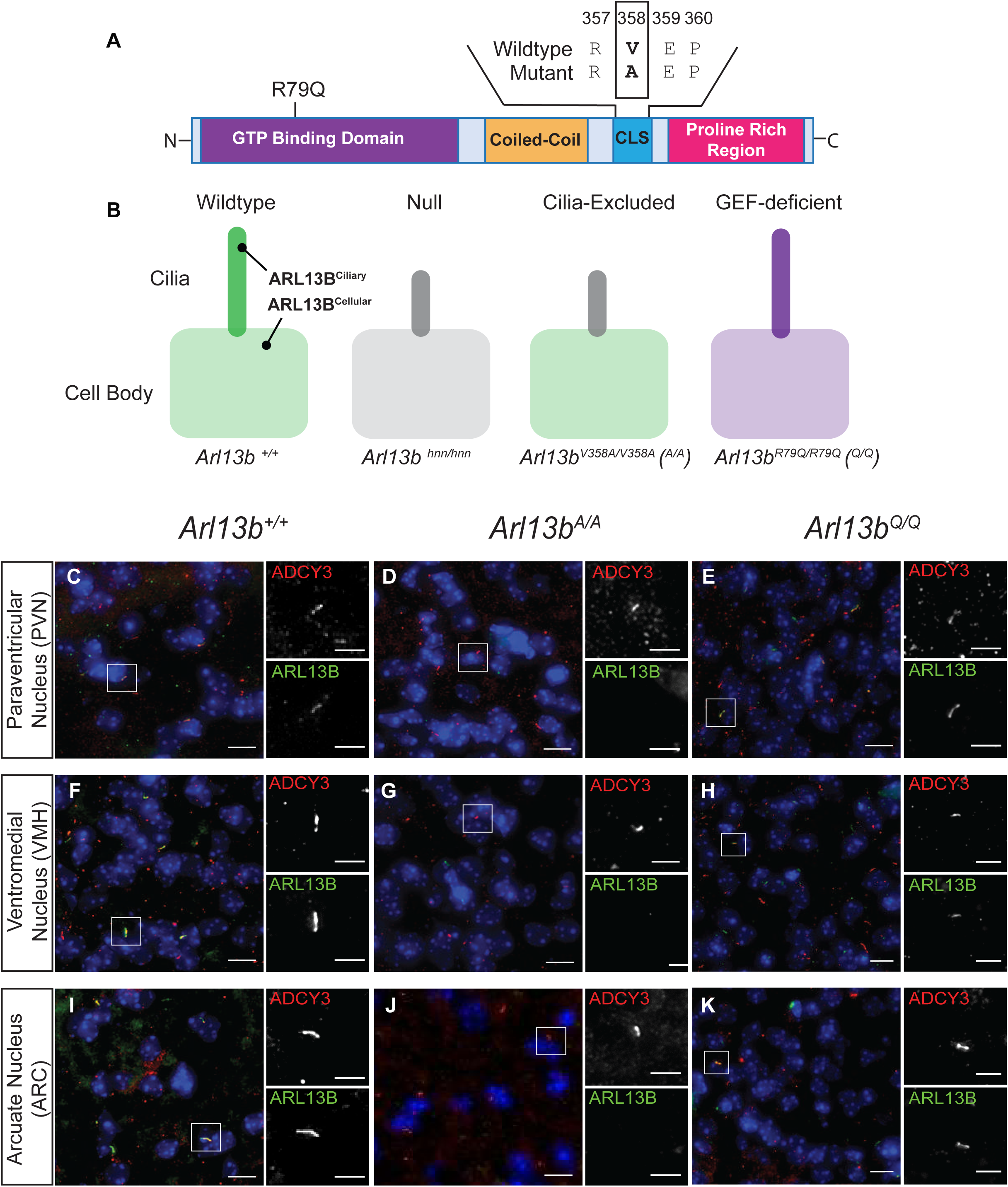
ARL13B^V358A^ protein is not detected in primary cilia of mouse tissues. (**A**) Schematic of ARL13B protein domains indicating the R79Q mutation in the GTP binding domain and amino acid sequence in wildtype and mutant in the cilia localization sequence (CLS). (**B**) Schematic of ARL13B localization in a wildtype (green, *Arl13b^+/+^*), null (gray, *Arl13b^hnn/hnn^*), cilia-excluded (*Arl13b^A/A^*), and GEF-deficient (purple, *Arl13b^R79Q/R79Q^*) models. (**C-K**) Immunofluorescence of ADCY3 (red) and ARL13B (green) in hypothalamic feeding centers, PVN (**C-E**), VMH (**F-H**), ARC (**I-K**) in the brain of postnatal day 7 *Arl13b^+/+^*, *Arl13b^A/A^*, and *Arl13b^Q/Q^* mice. Scale bars: 20μm and 5μm for insets indicated by white boxes. Hoechst-stained nuclei are blue.

In the pancreas, we performed immunofluorescence staining for ARL13B, acetylated α-tubulin (cilia), glucagon (α-cells), and insulin (β-cells). We observed ARL13B co-localizes with acetylated α-tubulin on α- and β-cells in the cilia of control mice but not on pancreatic islet cells in *Arl13b^A/A^* mice (**Supplemental Figure 2 A-D**). In the hypothalamus, we used the cilia marker adenylate cyclase 3 (ADCY3) to identify neuronal cilia (29). We detected ADCY3 staining in the paraventricular (PVN), the ventromedial (VMH), and the arcuate (ARC) nuclei, indicating the presence of neuronal cilia. In control mice, ARL13B co-localizes with ADCY3 in the PVN, VMH, and the ARC, indicating it is normally present in cilia (**Figure 3C**, **F**, **I**). However, we did not detect ARL13B in neuronal cilia in these brain regions of *Arl13b^A/A^* mice, indicating the engineered variant does not enrich in these cilia (**Figure 3D**, **G**, **J**). In addition to assessing cilia frequency, we measured the length of neuronal cilia in *Arl13b^A/A^* mice and found that the cilia length is shorter compared to *Arl13b^+/+^,* similar to what we previously reported in other cell types (**Supplemental Figure 3**) (22). These findings demonstrate that ciliary ARL13B is present in both the hypothalamic feeding centers and pancreatic islets in wildtype mice but is not detectable in cilia of mice expressing the ARL13B^V358A^variant.

### ARL13B’s GEF activity for ARL3 is unlikely to be required for body weight regulation

One possible mechanism of ARL13B action is via its GEF activity for ARL3. If activated ARL3 contributes to the regulation of energy homeostasis, then disrupting ARL13B’s ability to activate ARL3 should mimic the obesity phenotype observed in *Arl13b^A/A^* mice. The ARL13B^R79Q^ mutation abolishes ARL13B GEF activity for ARL3 *in vitro* without altering its cilia localization (14–16, 27, 30, 31) (**Figure 3B, E, H, K**). We found no significant difference in body weight between control and *Arl13b^R79Q/R79Q^* (*Arl13b^Q/Q^*) mice (**Figure 4A** and **B**). While we cannot exclude the possibility of a compensatory GEF for ARL3 in vivo, this finding suggests that the obesity phenotype in *Arl13b^A/A^* mice may be independent of ARL13B’s GEF activity.

**Figure 4:**
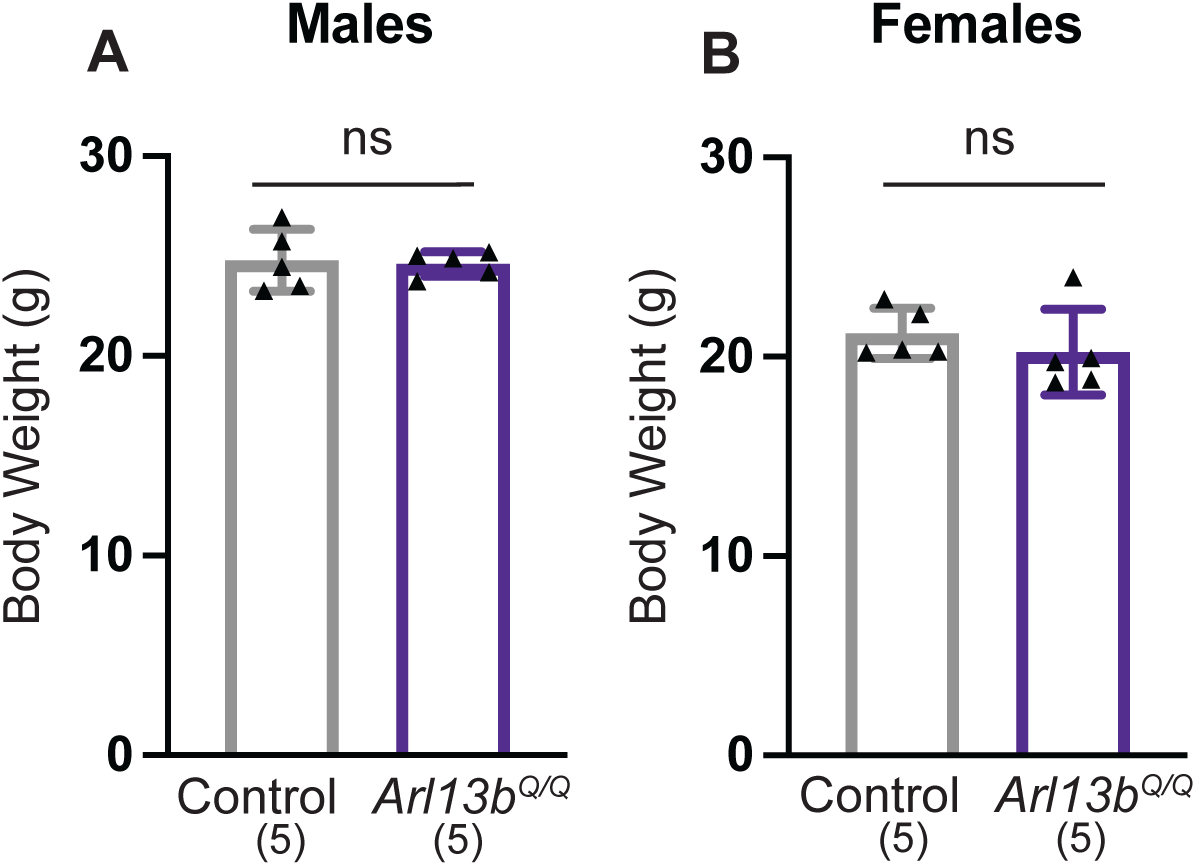
ARL13B’s GEF activity for ARL3 is not required for energy homeostasis. Body weights were assessed in adult (14–18-week-old) male (**A**) and female (**B**) Control (*Arl13b^+/+^*) and *Arl13b^Q/Q^* (*Arl13b^R79Q/R79Q^*) mice. Data are presented as means ± SD. Sample size indicated on graphs. ns, not significant with the Mann-Whitney U test, indicating *P* > 0.05.

### Ciliary ARL13B is required in cilia within the nervous system for control of body weight

A key question in defining the mechanism of ARL13B function within cilia is identifying the cell type in which ciliary ARL13B is required to regulate body weight. To genetically address this, we combined the *Arl13b^A^*allele with a conditional *Arl13b^floxed^* allele, which expresses wildtype ARL13B prior to recombination. This strategy allowed us to use tissue specific Cre recombination to delete the wildtype allele, leaving only the cilia excluded ARL13B^V358A^ protein (**Figure 5A**). Because our data indicate that ARL13B dosage does not contribute to the obesity phenotype (**Figure 1A and B**), this approach effectively isolates ARL13B function in a tissue specific and subcellular specific manner. We selected *nestin Cre* to target the nervous system, as primary cilia in neuronal populations are known to play critical roles in regulating body weight and food intake (3, 10, 11). *Nestin* expression begins in neural progenitors during midgestation, resulting in recombination within the nervous system (32). We found that both male and female *nestin-Cre;Arl13b^A/flox^* mice were significantly heavier than their control littermates (**Figure 5 B-C**). Together, these findings demonstrate that ciliary ARL13B function within the nervous system is essential for normal body weight regulation.

**Figure 5:**
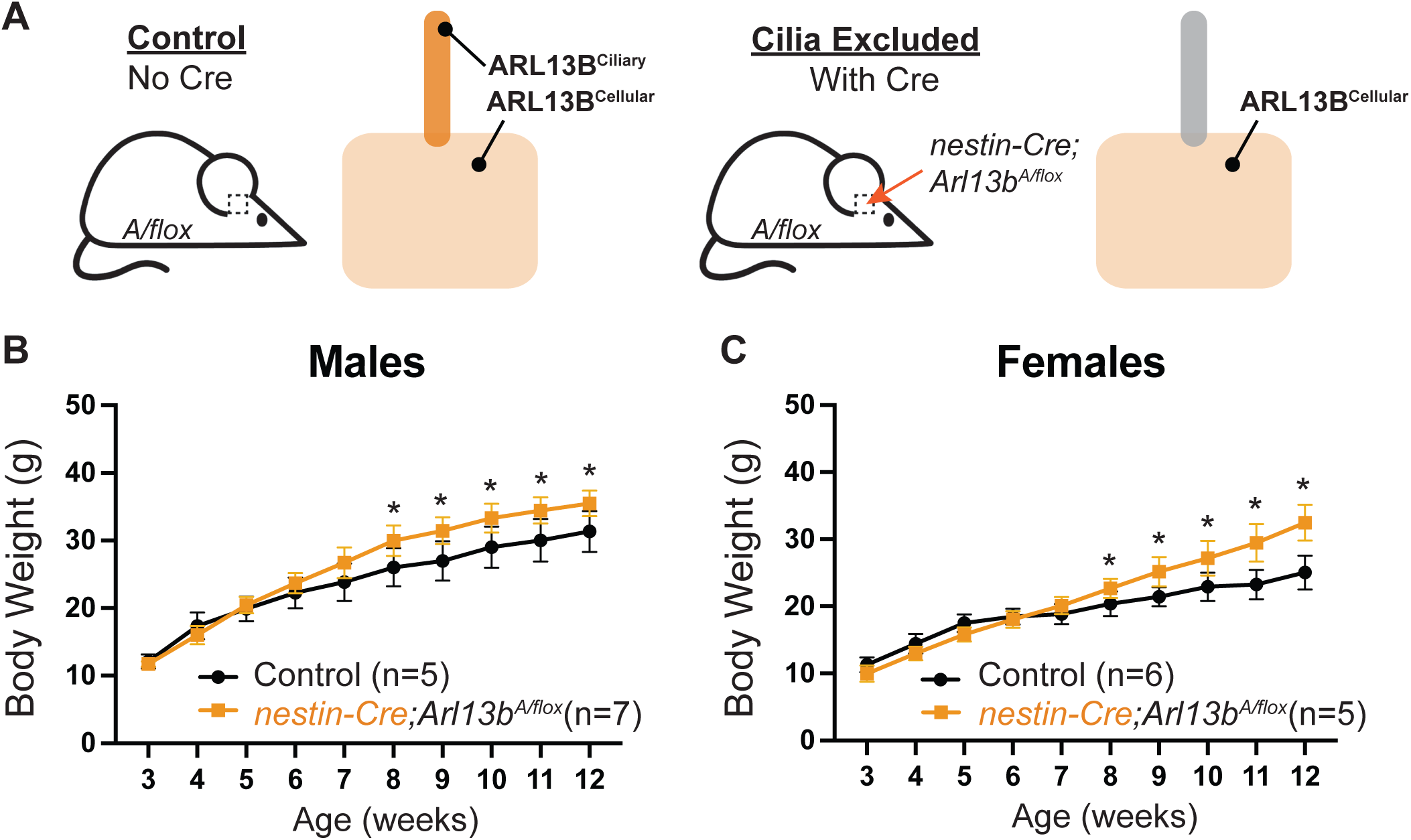
Ciliary ARL13B is required in nervous system cells to control body weight. **(A)** Schematic illustrating control mice (*Arl13b^A/flox^*) in which ARL13B localizes to both the cilium and cell, and cilia-excluded mice (*nestin-Cre;Arl13b^A/flox^*) in which ARL13B is selectively excluded from cilia in the nestin lineage cell population. (**B-C**) Weight curves of male (**B**) and female (**C**) control and *nestin-Cre;Arl13b^A/flox^* mice over time. Data are presented as means ± SD. Sample size indicated on graphs. Statistical significance was determined using repeated-measures ANOVA followed by Fisher’s LSD test. \**P* < 0.05.

### Rescuing ciliary ARL13B in 4-week-old Arl13b^A/A^ mice prevents obesity

Loss of ciliary ARL13B could control energy homeostasis through developmental defects or post-developmentally. To address this, we induced expression of wildtype ARL13B, which localizes to cilia, after development and tracked the body weight profile of the animals. We used a Cre-inducible *Arl13b-Fucci2a* (*AF2a*) allele that expresses wildtype ARL13B fused to a Cerulean fluorophore (33). We crossed *Arl13b^A/A^;AF2a* mice to *CAGG-CreER^TM^* transgenic mice in which Cre is ubiquitously activated following tamoxifen induction (**Figure 6A**). We confirmed that ARL13B-Cerulean is present in cilia only after Cre recombination (**Figure 6B** and **C**). ARL13B-Cerulean co-localizes with the neuronal marker ADCY3 in the PVN, the VMH, and the ARC in tamoxifen-treated *Arl13b^A/A^;AF2a;CAGG-CreER^TM^* mice but not in tamoxifen-treated *Arl13b^A/A^* mice lacking Cre (**Supplemental Figure 4 A-F**).

**Figure 6:**
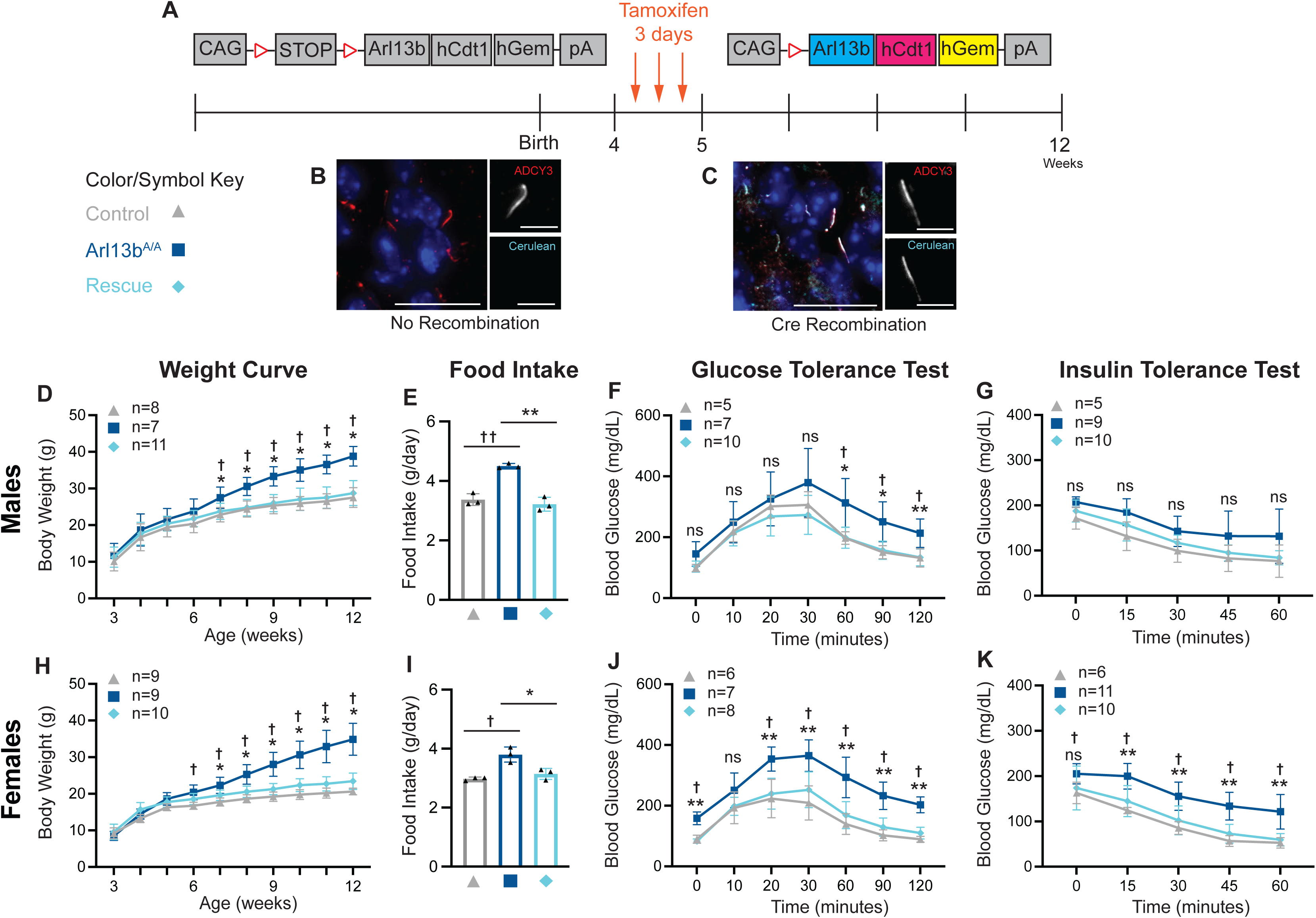
Rescuing ciliary expression of ARL13B prevents the metabolic phenotype of *Arl13b^A/A^* mice. (**A**) Schematic of the *Arl13b-Fucci2a (AF2a)* conditional allele with tamoxifen Cre induction timeline. (**B-C**) Immunofluorescence of ADCY3 (red) and ARL13B-Cerulean (cyan). (**B**) without Cre recombination, (**C**) with Cre-recombination. (**D and H**) Longitudinal body weight data of Control, Arl*13b^A/A^*, and Rescue (*Arl13b^A/A^;AF2a; CAGG-CreER^T2^*). (**E and I**) Food intake (performed in mice 6-7 weeks old). Control (males n=6; females n=7), *Arl13b^A/A^* (males n=7; females n=6) and Rescue (males n=7; females n=6). (**F and J**) Glucose tolerance test in males and females. Blood glucose was measured at the indicated times after i.p. glucose injection. (**G and K**) Insulin tolerance test in males and females. Blood glucose was measured at indicated times after i.p. insulin injection. Statistical significance was determined using repeated-measures ANOVA followed by Tukey’s multiple comparisons test. Asterisk (*) indicates *Arl13b^A/A^*is significantly different (*P* < 0.05) from Rescue; dagger (†) indicates *Arl13b^A/A^* is significantly different (*P* < 0.05) from control. \**P* < 0.05; \*\**P* < 0.01; †† *P* < 0.01; ns, not significant with *P* > 0.05. Data are presented as means ± SD. Sample size indicated on graphs.

We longitudinally tracked the impact of introducing ARL13B-Cerulean expression at 4-weeks in *Arl13b^A/A^;AF2a;CAGG-CreER^TM^* mice (referred to hereafter as *Arl13b^A/A^;AF2a^CAGG-4wk^*) compared to controls *Arl13b^A/+^;AF2a;CAGG-CreER^TM^* and *Arl13b^A/A^* mice. From week 7 until week 12, both male and female *Arl13b^A/A^; AF2a^CAGG-4wk^* mice weighed a similar amount to the control mice and significantly less than *Arl13b^A/A^* mice (**Figure 6D** and **H**). These data demonstrate that ciliary ARL13B plays a role after development is complete.

To determine whether the reduction in body weight is the result of changes in feeding, we measured food intake two weeks post-tamoxifen administration, just prior to when we observed the differentiation in body weight between *Arl13b^A/A^ and Arl13b^A/A^;AF2a^CAGG-4wk^*mice. We found that male and female *Arl13b^A/A^;AF2a^CAGG-4wk^*mice consumed significantly less food per day compared to *Arl13b^A/A^* (**Figure 6E** and **I**).

To examine the homeostatic response of *Arl13b^A/A^;AF2a^CAGG-4wk^*mice to glucose and insulin, we performed glucose and insulin tolerance tests in 13 to 15-week-old mice. In response to glucose, male and female *Arl13b^A/A^;AF2a^CAGG-4wk^*mice cleared glucose from their bloodstream, similar to control mice, while *Arl13b^A/A^* mice displayed elevated blood glucose levels (**Figure 6F** and **J**). In response to insulin, blood glucose levels in male *Arl13b^A/A^;AF2a^CAGG-4wk^* mice trended toward the level in control animals (**Figure 6G**). In contrast, insulin lowered blood glucose levels in female *Arl13b^A/A^;AF2a^CAGG-4wk^*mice (**Figure 6K**). These data demonstrate that introducing wildtype ARL13B in cilia of *Arl13b^A/A^*mice normalizes glycemic regulation, showing that ARL13B must be in cilia for normal glucose and insulin metabolism. Taken together, the restoration of body weight as well as food intake and homeostatic response in *Arl13b^A/A^;AF2a^CAGG-4wk^*mice is consistent with ciliary ARL13B controlling acute signaling.

To more specifically evaluate the role of ciliary ARL13B in hypothalamic development, we assessed the integrity of hypothalamic circuits that control feeding behavior. Appetite-stimulating agouti-related peptide (AgRP)-expressing neurons and the appetite-suppressing pro-opiomelanocortin (POMC)-expressing neurons of the ARC are key regulators of food intake (34). To assess whether loss of ciliary ARL13B disrupted the number, patterning, or projections of AgRP and POMC neurons in the ARC, we performed RNAscope *in situ* hybridization and immunofluorescence analysis. We found normal numbers and patterning for both AgRP and POMC neurons in *Arl13b^A/A^* mice (**Supplemental Figures 5 A-D**). Similar to what we previously reported in other cell types, we observed no significant changes in AgRP and POMC neuron projections in *Arl13b^A/A^* mice (**Supplemental Figures 5 E-G**) (23, 35). These findings show that the absence of ciliary ARL13B during development does not impair the number, patterning, or projections of AgRP or POMC neurons in the ARC and suggest a potential role for ciliary ARL13B in adult homeostasis. In light of the data showing the rescue of body weight, food intake, and homeostatic responses in *Arl13b^A/A^;AF2a^CAGG-4wk^*mice, these data support a model in which ciliary ARL13B plays a post-developmental role.

## DISCUSSION

Our findings define a cilia-specific role for ARL13B as an essential regulator of energy homeostasis. Mice homozygous for the cilia-excluded variant, ARL13B^V358A^, became hyperphagic, obese, and insulin resistant shortly after weaning. Using a genetic toolkit, we showed that selective loss of ciliary ARL13B in the nervous system caused obesity, while post-developmental restoration of ciliary ARL13B fully rescued the feeding behavior and obesity phenotypes. Furthermore, our analysis showed no overt changes in cell population numbers or projections within the ARC. These data argue against a developmental requirement and instead suggest that ciliary ARL13B is acutely required to maintain signaling pathways that govern energy balance in adult mice. These findings advance the field, demonstrating that the ciliary function of a protein, independent of its non ciliary roles, is required for energy homeostasis. In contrast, prior studies have relied on complete cilia ablation (e.g., via conditional IFT mutations) or on mutations that disrupt both ciliary and cellular pools of signaling proteins, such as GPCRs (1).

While ARL13B^V358A^ is an engineered mutation, we were surprised to observe obesity in the engineered *Arl13b^A/A^* mice since distinct patient mutations in *ARL13B* lead to Joubert syndrome (JS) (20, 21). JS is clinically defined by its neurodevelopmental features and does not typically include obesity as a feature. Consistent with this, mice expressing the JS-causing variant, ARL13B^R79Q^ do not exhibit a weight phenotype. However, there are clear links between cilia and obesity (36). These include ciliopathies like BBS and ALMS or mutations in the G-protein-coupled receptor melanocortin receptor 4 (MC4R), which localizes to cilia (37, 38). Thus, the cilia-specific role of ARL13B in controlling energy homeostasis may involve interactions with other ciliopathy proteins or signaling machinery.

Our data clearly argue that obesity in *Arl13b^A/A^* animals is driven by hyperphagia and increased fat mass. We eliminated two alternative possibilities for the weight phenotype. First, we previously reported that *Arl13b^A/A^* mice develop progressive, mild kidney cysts (28). However, the size and rate of kidney cyst progression in these animals do not account for the changes in body weight and composition we observe (28). Second, we showed that the weight phenotype is independent of gene dosage, as *Arl13b^A/hnn^* mice displayed a body weight phenotype similar to *Arl13b^A/A^* mice. Similarly, the expression of one or two copies of the *Arl13b-Cerulean* allele in *Arl13b^+/+^*, *Arl13b^hnn/+^*, and *Arl13b^hnn/hnn^* mice did not display a weight phenotype (39).

To evaluate whether the hyperphagia was driven by the nervous system (**Figure 1A and B**), we generated *nestin-Cre;Arl13b^A/flox^* mice. Although the weight phenotype in *nestin-Cre;Arl13b^A/flox^* mice was less severe than that observed in *Arl13b^A/A^* mice, this difference may be influenced by the *nestin Cre* allele itself, as hemizygous *nestin Cre* mice exhibit mild growth retardation (40). Importantly, despite this potential modulation, *nestin-Cre;Arl13b^A/flox^* mice were significantly heavier than their control littermates, implicating cells within the nervous system as critical mediators of ARL13B’s effects on energy homeostasis. More critically, these results indicate that this approach successfully isolates cell populations in which ciliary ARL13B is required for body weight control and motivate future studies to define the specific neuronal populations involved, which should provide important insights into the ciliary mechanism of ARL13B action.

While our data argue that obesity in *Arl13b^A/A^*animals is driven by hyperphagia controlled by neuronal cell populations, other studies have shown that ciliary dysfunction in peripheral tissues, including adipose tissue and the pancreas, is linked to metabolic imbalance (41). Thus, it is important to acknowledge that ciliary ARL13B from the periphery could contribute in some way to the *Arl13b^A/A^* phenotype. For example, inhibition of BBS12 expression in human adipocyte precursor cells increases the expression of key regulators of adipogenesis and promotes adipose tissue expansion in *Bbs12^-/-^*mice (42). In the pancreas, loss of primary cilia in β cells disrupts insulin secretion and leads to impaired glucose homeostasis (26). Chronically elevated insulin levels promote lipogenesis and contribute to the development of metabolic disease. Future studies will need to examine the contribution of ciliary ARL13B to obesity-related phenotypes in these peripheral cell types.

One possible mechanism through which ciliary ARL13B regulates energy homeostasis involves the established interaction between ARL13B and INPP5E in the ciliary membrane (16, 17, 30). Patients lacking the C-terminal CaaX domain of INPP5E lose INPP5E ciliary localization and exhibit MORM syndrome, whose symptoms include obesity (43). INPP5E is not detected in the cilia of *Arl13b^A/A^* cells in culture, consistent with data showing that the ciliary retention of INPP5E depends on ARL13B (22). Thus, the loss of ciliary INPP5E in the absence of ciliary ARL13B could impact the trafficking or signaling of proteins implicated in feeding behavior, such as MC4R. This would suggest the obesity-causing ARL13B^V358A^ mutation functions via INPP5E, whereas JS-causing mutations in ARL13B act via ARL3. While a tantalizing model, it is challenging to reconcile with *INPP5E* mutations causing JS or the finding that INPP5E requires activated ARL3 (ARL3-GTP) to enter cilia.

Our *Arl13b^A/A^* model isolates the ciliary role of the ARL13B protein, enabling direct interrogation, at subcellular resolution, of ARL13B controlling cilia-mediated signaling pathways involved in maintaining energy homeostasis. Our ability to combine subcellular resolution with tissue specificity holds deep promise for future studies to identify the cell type or combination of cell types that require ciliary ARL13B function. Thus, we provide the field an important set of models from which to resolve the mechanism of ciliary ARL13B action to regulate energy homeostasis.

## Acknowledgements

The authors would like to acknowledge the Mouse Metabolism Core at the UCSF Nutrition & Obesity Center (NIH NIDDK P30DK098722) for technical support and assistance with metabolic phenotyping experiments. The authors also acknowledge the staff within the Parnassus Advanced Light Microscopy CoLab (PALM) at UCSF Parnassus Heights for their training and support in using Imaris 9.5.1; and the Center for Advanced Light Microscopy (CALM) at UCSF for their training and support in using the CSU wide field confocal microscope (obtained using the NIH S10 Shared Instrumentation Grant: 1S10OD017993-01A1).

## Author contributions

Conceptualization TTT, TC, EDG. Investigation EDG, TTT, CMA, KMB, XY. Formal analysis TTT, CMA. Visualization TTT. Funding acquisition TTT, TC, NFB, CV, XY, EDG, CMA, KMB

Writing initial draft. TTT. Writing, reviewing, and editing TTT, TC, NFB, CV, KMB, XY. Project administration TC, CV, NFB.

## Guarantor Data Access and Responsibility

Dr. Caspary is the guarantor of this work, had full access to all the data, and takes full responsibility for the integrity of data and the accuracy of data analysis.

## Prior Presentation

Parts of this study were presented at the Biology of Cilia and Flagella Conference 2024, Minnesota, USA, May 12-16, and the ASCB Cell Bio 2025 Conference, Pennsylvania, USA, December 7.

## Supplemental Figure Legends

**Figure S1: Metabolic Analysis.**

Metabolic analysis was measured over 48 hours using indirect calorimetry and CalR version 2. Control (*Arl13b^A/+^, Arl13b ^+/+^*) mice (males n=10; females n=9; dark grey) and *Arl13b^A/A^* mice (males n=8, blue; females n=9, orange). (**A, H**) Energy expenditure was plotted as a function of lean mass for each group. Lines represent group regression, and each point represents one mouse. (**A-C, H-J**) Energy Expenditure. (**D, E, K, L**) Locomotor activity. (**F, G, M, N**) Respiratory exchange ratio (RER). Grey areas indicate the dark photoperiod. (**B, D, F, I, K, M**) Time-course data are shown as mean ± SEM for each genotype (bold line and shaded area). Values are plotted hourly. (**C, E, G, J, L, N**) Boxplots showing the 5th percentile, median, and 95th percentile of total (48 h), dark-phases, and light-phases. Each point represents one mouse. Homogeneity of slopes was assessed using a general linear model (GLM) on GraphPad. Genotype differences were analyzed by ANCOVA with lean mass (EchoMRI, 6-week-old mice) as a covariate, except in panel J, where a significant interaction necessitated a GLM with lean mass*Group.

**Figure S2: ARL13B^V358A^ protein is not detected in α-cell and β-cell cilia in the pancreas.**

(**A-D**) Immunofluorescence staining of acetylated α-tubulin (white –“AcTub”), ARL13B (green), glucagon and insulin (red) in α-cells (**A-B**) and β-cells (**C-D**) in the mouse pancreas. Hoechst-stained nuclei are blue. Scale bars: 20μm and 5μm for insets.

**Figure S3: Neuronal cilia length analysis in *Arl13b^A/A^* mice.**

Quantification of neuronal cilia length in *Arl13b^+/+^*and *Arl13b^A/A^* mice. Data points represent individual cilia. Data are presented as means ± SD. Statistical significance was determined using the Mann-Whitney U test. ** *P* < 0.01.

**Figure S4: ARL13B-cerulean protein is detectable in cilia in adult mouse tissues.**

(**A-F**) Representative immunofluorescence staining of ADCY3 (red) and ARL13B-Cerulean (cyan) in hypothalamic feeding centers, PVN (**A-B**), VMH (**C-D**), ARC (**E-F**), in brain tissue from tamoxifen-treated *Arl13b^A/A^* and *Arl13b^A/A^; AF2a^CAGG-4wk^* mice at 12 weeks of age. Scale bars: 20μm and 5μm for insets indicated by white boxes. Hoechst-stained nuclei are blue.

**Figure S5: Loss of ciliary ARL13B does not disrupt neuronal patterning or axonal projections of AgRP or POMC neurons.**

(**A-B**) Images of RNAscope *in situ* hybridization of AgRP (red) and POMC (blue) neurons in the ARC of control (left column) and *Arl13b^A/A^* (right column) mice. (**C-D**) Quantification of AgRP (**C**) or POMC (**D**) neurons in each genotype. Data are presented as means ± SD. At least 3 animals per group. ns, not significant, with Student’s T-test indicating *P* > 0.05. (**E**) Representative immunofluorescence images of AgRP (green, upper row) and αMSH (magenta, lower row) in the PVN (outlined in white dotted line) of control (left column) or *Arl13b^A/A^* (right column) mice. (**F-G**) Superplot of AgRP (**F**) or POMC (**G**) neuron axon density in the PVN. Each circle represents the axon density of one PVN section (3 per mouse). Each triangle represents the axon density of one mouse (n ≥ 3 animals per group). Data are presented as mouse means ± SD. (AgRP: p=0.0653, POMC: p=0.4910). ns, not significant with Welch’s t-test conducted on axon density per mouse. Scale bar represents 100 µm.

