## Supplementary material for "Ciliary ARL13B is essential for body weight regulation in mice": Supplmental Materials

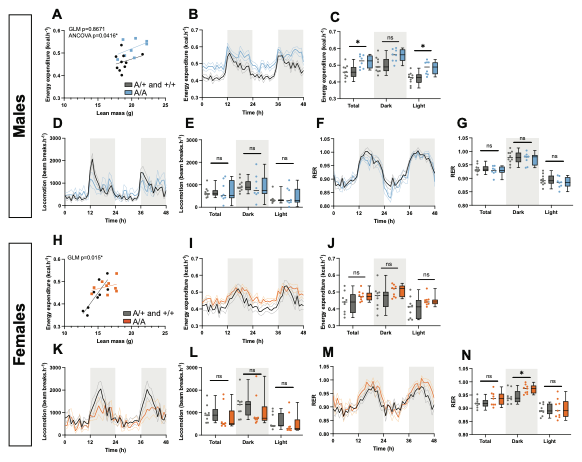

**Figure S1**: **Metabolic Analysis.**

Metabolic analysis was measured over 48 hours using indirect calorimetry and CalR version 2. Control (*Arl13b^A/+^, Arl13b ^+/+^*) mice (males n=10; females n=9; dark grey) and *Arl13b^A/A^* mice (males n=8, blue; females n=9, orange). (**A, H**) Energy expenditure was plotted as a function of lean mass for each group. Lines represent group regression, and each point represents one mouse. (**A-C, H-J**) Energy Expenditure. (**D, E, K, L**) Locomotor activity. (**F, G, M, N**) Respiratory exchange ratio (RER). Grey areas indicate the dark photoperiod. (**B, D, F, I, K, M**) Time-course data are shown as mean ± SEM for each genotype (bold line and shaded area). Values are plotted hourly. (**C, E, G, J, L, N**) Boxplots showing the 5th percentile, median, and 95th percentile of total (48 h), dark-phases, and light-phases. Each point represents one mouse. Homogeneity of slopes was assessed using a general linear model (GLM) on GraphPad. Genotype differences were analyzed by ANCOVA with lean mass (EchoMRI, 6-week-old mice) as a covariate, except in panel J, where a significant interaction necessitated a GLM with lean mass*Group.

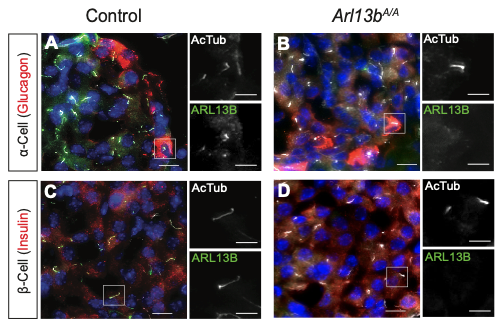

**Figure S2**: **ARL13B^V358A^ protein is not detected in α-cell and β-cell cilia in the pancreas.**

(**A-D**) Immunofluorescence staining of acetylated α-tubulin (white –“AcTub”), ARL13B (green),

glucagon and insulin (red) in α-cells (**A-B**) and β-cells (**C-D**) in the mouse pancreas.

Hoechst-stained nuclei are blue. Scale bars: 20μm and 5μm for insets.

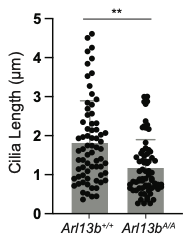

**Figure S3**: **Neuronal cilia length analysis in *Arl13b^A/A^* mice.**

Quantification of neuronal cilia length in *Arl13b^+/+^* and *Arl13b^A/A^* mice. Data points represent individual cilia. Data are presented as means ± SD. Statistical significance was determined using the Mann-Whitney U test. ** *P* < 0.01.

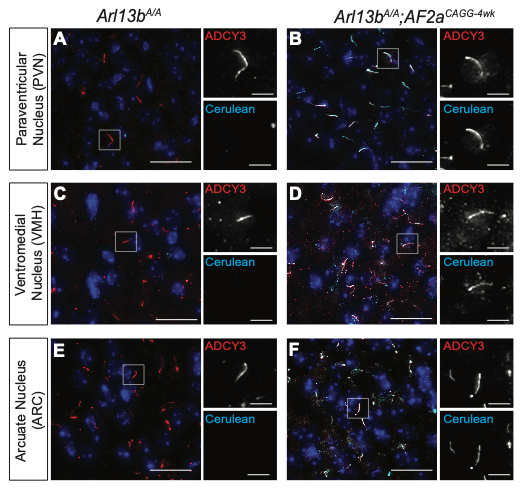

**Figure S4**: **ARL13B-cerulean protein is detectable in cilia in adult mouse tissues.**

(**A-F**) Representative immunofluorescence staining of ADCY3 (red) and ARL13B-Cerulean (cyan) in hypothalamic feeding centers, PVN (**A-B**), VMH (**C-D**), ARC (**E-F**), in brain tissue from tamoxifen-treated *Arl13b^A/A^* and *Arl13b^A/A^; AF2a^CAGG-4wk^* mice at 12 weeks of age. Scale bars: 20μm and 5μm for insets indicated by white boxes. Hoechst-stained nuclei are blue.

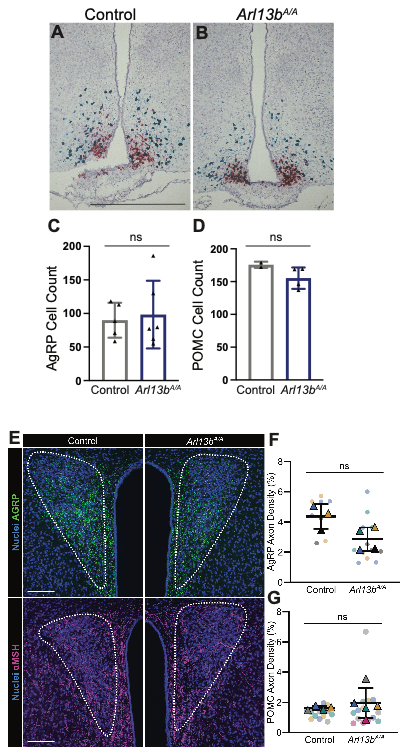

**Figure S5**: **Loss of ciliary ARL13B does not disrupt neuronal patterning or axonal projections of AgRP or POMC neurons.**

(**A-B**) Images of RNAscope *in situ* hybridization of AgRP (red) and POMC (blue)

neurons in the ARC of control (left column) and *Arl13b^A/A^* (right column) mice. (**C-D**) Quantification of AgRP (**C**) or POMC (**D**) neurons in each genotype. Data are presented as means ± SD. At least 3 animals per group. ns, not significant, with Student’s T-test indicating *P* > 0.05. (**E**) Representative immunofluorescence images of AgRP (green, upper row) and αMSH (magenta, lower row) in the PVN (outlined in white dotted line) of control (left column) or *Arl13b^A/A^* (right column) mice. (**F-G**) Superplot of AgRP (**F**) or POMC (**G**) neuron axon density in the PVN. Each circle represents the axon density of one PVN section (3 per mouse). Each triangle represents the axon density of one mouse (n ≥ 3 animals per group). Data are presented as mouse means ± SD. (AgRP: p=0.0653, POMC: p=0.4910). ns, not significant with Welch’s t-test conducted on axon density per mouse. Scale bar represents 100 µm.
